# Transferrin binding protein B and Transferrin binding protein A 2 expand the transferrin recognition range of *Histophilus somni*

**DOI:** 10.1101/730739

**Authors:** Anastassia K. Pogoutse, Trevor F. Moraes

**Affiliations:** Department of Biochemistry, University of Toronto, Toronto, Ontario, Canada

**Keywords:** *Histophilus somni*, Lipoproteins, Membrane Transport Proteins, Transferrins, Transferrin binding protein A, Transferrin binding protein B

## Abstract

The bacterial bipartite transferrin receptor is an iron acquisition system that is required for survival by several key human and animal pathogens. It consists of the TonB-dependent transporter Transferrin binding protein A (TbpA) and the surface lipoprotein Transferrin binding protein B (TbpB). Curiously, the Tbps are only found in host specific pathogens, and are themselves host specific, meaning that they will bind to the transferrin of their host species, but not to those of other animal species. While this phenomenon has long been established, neither the steps in the evolutionary process that led to this exquisite adaptation for the host, nor the steps that could alter it, are known. We sought to gain insight into these processes by studying Tbp specificity in *Histophilus somni*, a major pathogen of cattle. A past study showed that whole cells of *H. somni* specifically bind bovine transferrin, but not transferrin from sheep and goats, two bovids whose transferrins share 93% amino acid sequence identity with bovine transferrin. To our surprise, we found that *H. somni* can use sheep and goat transferrins as iron sources for growth, and that *Hs*TbpB, but not *Hs*TbpA, has detectable affinity for sheep and goat transferrins. Furthermore, a third transferrin binding protein, *Hs*TbpA2, also showed affinity for sheep and goat transferrins. Our results show that *H. somni* TbpB and TbpA2 act to broaden the host transferrin recognition range of *H. somni*.

**Importance:** Host restricted pathogens infect a single host species or a narrow range of host species. *Histophilus* somni, a pathogen that incurs severe economic losses for the cattle industry, infects cattle, sheep, and goats, but not other mammals. The transferrin binding proteins, TbpA and TbpB, are thought to be a key iron acquisition system in *H. somni*, however, surprisingly, they were also shown to be cattle transferrin-specific. In our study we find that *H. somni* TbpB, and another little-studied Tbp, TbpA2, bind sheep and goat transferrins as well as bovine transferrin. Our results suggest that TbpA2 may have allowed for host range expansion, and provide a mechanism for how host specificity in Tbp containing pathogens can be altered.

## Introduction

Virulence factor specificity arises over the course of evolutionary arms races between pathogens and their hosts (1–4). In this conflict, mutations that help the host evade the pathogen confer a fitness advantage, and in turn, compensatory mutations that allow the pathogen to continue exploiting the host confer an advantage to the pathogen. After multiple rounds of coevolution, the pathogen becomes uniquely adapted to the host and may lose affinity for proteins found in other host species, resulting in specificity (5). Host-species-specific interactions have been identified in a virulence factor known as the bacterial bipartite transferrin receptor (6). This receptor is found in some members of the Beta- and Gammaproteobacteria and consists of two proteins: the TonB-dependent transporter Transferrin binding protein A (TbpA) and the surface lipoprotein (SLP) Transferrin binding protein B (TbpB). The Tbps are employed by bacteria to commit a type of iron piracy, in which iron is removed from the C-terminal lobe of transferrin and transported across the bacterial outer membrane (7). A TbpA-driven evolutionary arms race between primate pathogens and their hosts has left signatures in the transferrins of some primates (including humans) (8).

Both TbpA and TbpB can bind transferrin independently of each other. Because the binding surface of TbpB extends further out from the membrane, TbpB is thought to fish out transferrin and pass it to TbpA. TbpA removes the iron from the transferrin’s C-lobe and transports the Fe^3+^ across the outer membrane (9, 10); the transferrin is then released from the receptor. TbpA performs the essential transport function while TbpB increases the efficiency of the receptor by preferentially binding holo-transferrin (11). While only TbpA acts as a transporter, both receptor proteins have been shown to be essential for infection (12).

Both TbpA and TbpB selectively bind the transferrins of their hosts. Furthermore, the specificity of the Tbp-transferrin interaction is a significant contributor to the host restriction observed in these pathogens. This has been observed in mouse models of neisserial infection and has caused some researchers to use transgenic transferrin mice in their experiments (13, 14). Understanding which subset of intermolecular contacts restricts the range of ligands a protein can bind could lend insight into the Tbp mechanism of function, as well allow for the design of better animal models.

Apart from TbpA and TbpB, another notable transferrin receptor is Transferrin binding protein A 2. TbpA2 is named based on its resemblance to TbpA – both are TonB-dependent transporters that bind transferrin, although TbpA2 is smaller than TbpA (85 kDa compared to 100 kDa). It has also been shown to bind the N-lobe of transferrin rather than the C-lobe (15). To date, TbpA2 has only been identified in two species, the livestock pathogens *Pasteurella multocida* and *Histophilus somni. P. multocida* does not encode a TbpBA system while *H. somni* does. In the latter organism, TbpA2 is thought to undergo phase variable expression (16). Little is known about TbpA2’s mechanism of function.

Out of all Tbp-encoding bacteria, *H. somni* presents a particularly interesting system for studying transferrin binding specificity. *H. somni* is a Gram-negative host-restricted opportunistic pathogen in the *Pasteurellaceae* family. In cattle, *H. somni* has been associated with bronchopneumonia, myocarditis, infectious thrombotic meningoencephalitis, arthritis, mastitis, and spontaneous abortion (17–22). It has also been associated with disease in domestic sheep, bighorn sheep, and bison (20, 22, 23), and has been isolated from the nasal and reproductive tracts of goats (24, 25). It is unclear whether strains of *H. somni* that circulate in one animal species can be transmitted to other species (23). In one study, cattle (bovine) strains of *H. somni* were shown to bind bovine transferrin, but not sheep (ovine), goat (caprine), or human transferrin (15, 26).The Tbp system is likely to be the receptor *H. somni* uses to take up iron during colonization, as the targets of its other iron receptors, such as heme or hemoglobin, are normally intracellular. Also, the Tbps are generally thought to be important in *H. somni* virulence based on the finding that bovine transferrin increases morbidity and mortality in a mouse model of *H. somni* infection (27). Therefore, the finding that bovine isolates of *H. somni* selectively bind bovine transferrin suggested that different strains of *H. somni* were specialized to infect different species of ruminants, with some being cattle-specific. Because the percent amino acid sequence identity between bovine and ovine, and bovine and caprine transferrins is 93%, we determined that relatively few intermolecular contacts in these interfaces must be specificity determinants in the *Hs*Tbp-transferrin interaction. This ability to differentiate between closely-related transferrins suggests that relatively few mutations would be required to determine which intermolecular contacts drive specific recognition by the *H. somni* Tbps. However, we first sought to reproduce the previously published results and to determine whether each TbpA, TbpB, and TbpA2 each displayed the same level of selectivity as was observed in the whole cell binding assay.

We selected strain H191, a bovine isolate of *H. somni*, for our investigations into Tbp binding specificity. Contrary to prior observations, we found that whole cells of our *H. somni* bovine isolate bind both bovine and ovine transferrin (although the cells bound ovine transferrin with a relatively lower affinity) (28). Under some conditions, *H. somni* was also able to use both ovine and caprine transferrins as iron sourcesfor growth. By overexpressing TbpA and TbpB separately in *E. coli*, we determined that while this strain’s TbpA displays no observable affinity for ovine or caprine transferrin, the TbpB binds ovine and caprine transferrins with a sub-micromolar affinity. Furthermore, we also identified TbpA2 in strain H191, expressed it in *E. coli*, and determined that, while it preferentially binds bovine transferrin, it also binds ovine and caprine transferrins. Taken together, our results suggest that *H. somni* TbpB and TbpA2 may act to broaden the host transferrin recognition range of this pathogen.

## Results

### *H. somni* preferentially takes up iron from bovine transferrin, but also shows affinity for ovine and caprine transferrins

In order to see if we could recapitulate the results of the previously published study, we performed a binding experiment with whole fixed cells of *H. somni* strain H191. We performed a competitive assay known as a displacement ELISA, and looked at binding of whole fixed cells of *H. somni* to bovine, ovine, caprine, and porcine transferrins. Cattle, sheep, and goats are known hosts of *H. somni* while swine are not, and so porcine transferrin was used as a negative control (Figure 1A). *H. somni* bound bovine transferrin, as expected, but also displayed some affinity for ovine transferrin. There was no statistically significant binding to the caprine or porcine transferrins (Figure 1B).

**Figure 1.**
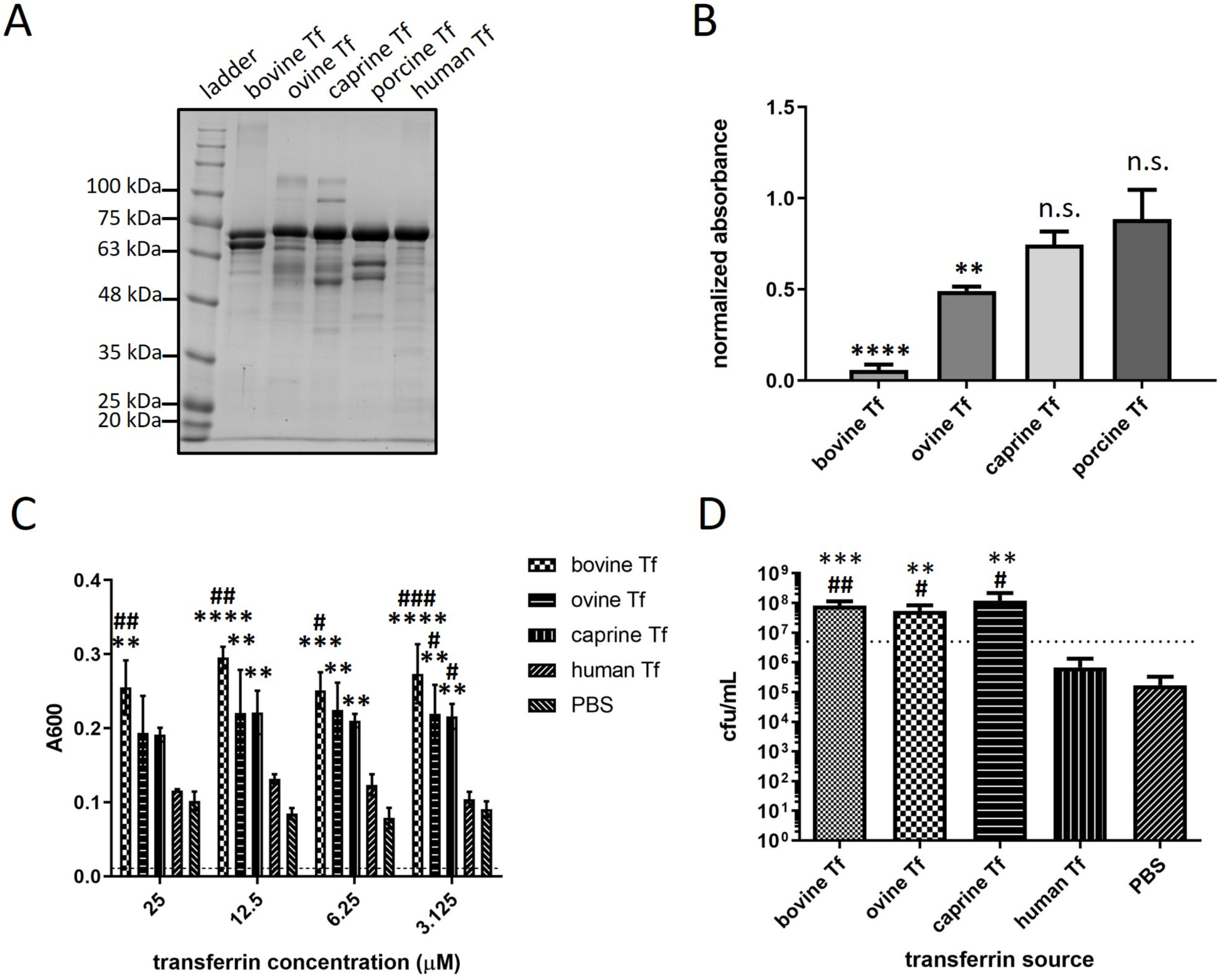
Whole-cell ELISAs and growth assays with *Histophilus somni* measuring binding to and growth on various transferrins. (A) Purified transferrins used in this study. (B) Binding of *H. somni* to various transferrins (bovine, ovine, caprine, and porcine). Whole cells of *H. somni* str. H191 were fixed with paraformaldehyde and used in a displacement ELISA. A 20-fold excess of unlabeled competitor transferrin was mixed with biotinylated bovine transferrin and incubated with the cells. Binding was detected using streptavidin-HRP and a colorimetric substrate (see Methods, section 4.3.3). All signal was normalized to “no competitor” absorbance readings. Data was analyzed using a one-way ANOVA followed by Dunnett’s test, with all values compared to the “no competitor” column. (C) Growth of *H. somni* in the presence of various transferrins (bovine, ovine, caprine, and human) at 25, 12.5, 6.25, and 3.125 µM as well as buffer alone. (D) Growth of *H. somni* in the presence of the same transferrin iron sources as in (C), at 25 µM following a “pre-starvation” step (see Methods 4.3.10). Dashed lines indicate cell density at the time of inoculation. Asterisks indicate levels of growth that were significantly higher than on buffer alone. Hashtags indicate levels of growth that were significantly higher than growth on human transferrin. Results were analyzed using a one-way ANOVA followed by Dunnett’s multiple comparison test. * indicates p<0.05, ** indicates p<0.01, *** indicates p<0.001, **** indicates p<0.0001. All values represent the mean of 3 separate experiments, and error bars represent standard error of the mean.

Next, we asked whether *H. somni* selectively uses bovine transferrin for growth, or whether its binding of ovine transferrin leads to iron uptake from both of these iron sources. We measured growth in the presence of bovine, ovine, caprine, and human transferrins at various concentrations, as well as in buffer alone (Figure 1C). *H. somni* does not normally infect humans and so human transferrin was used as a negative control. *H. somni* grew significantly better in the presence of bovine transferrin than in the presence of human transferrin at every transferrin concentration. Minimal growth was observed in the presence of human transferrin and in the buffer-only condition. *H. somni* also showed growth on ovine and caprine transferrins at every concentration except 25 µM. Growth on ovine and caprine transferrins was significantly greater than growth on human transferrin only at the lowest concentration (Figure 1C). Overall, the results demonstrate that *H. somni* has a lower affinity for ovine and caprine transferrins than for bovine. However, it does appear that *H. somni* can utilize ovine and caprine transferrin.

We wondered if the relatively low levels of growth on ovine and caprine transferrin at 25 µM was due to a lower number of transferrin receptors being expressed at the cell surface under more stringent conditions. This low expression may be due to the sudden shift between an iron-replete and iron-starving environment. To address this, we performed another experiment where we “pre-starved” the cells by growing them under iron-limiting conditions that were not as stringent as those used in the previous experiment (see Methods). In this case, following the pre-starvation step, cells showed growth in the presence of bovine, ovine, and caprine transferrins at 25 µM when compared to growth on human transferrin or buffer alone (Figure 1D).

We concluded that *H. somni* H191 does not exhibit strict specificity for bovine transferrin. However, it does display a preference for bovine transferrin while also having affinity for transferrins of other bovids. To determine which transferrin binding protein(s) was responsible for this binding profile, we went on to investigate each Tbp separately in a heterologous system.

### *Hs*TbpA selectively binds bovine transferrin while *Hs*TbpB and *Hs*TbpA2 show greater promiscuity

To address the binding contribution of each of the transferrin binding proteins (TbpA, TbpA2 and TbpB), we expressed His-tagged TbpAs, TbpA2, and GST-tagged *Hs*TbpB independently in *E. coli*, and performed displacement ELISAs (Figure 2). *E. coli* does not encode any of these three receptors and so this served as an appropriate heterologous system. TbpA assays were performed with whole cells while TbpB assays were performed with the soluble lysate fraction (TbpB was expressed without its N-terminal cysteine and was therefore not lipidated). An interaction was denoted as “specific” when unlabeled bovine transferrin lowered the absorbance signal to 10% or less of the maximum and other unlabeled competitor transferrins did not decrease the signal significantly. TbpA specifically bound bovine transferrin (Figure 2B). Control TbpAs from the ruminant pathogen *Mannheimia haemolytica* and from the porcine pathogen *Actinobacillus pleuropneumoniae* were used to validate the assay. The TbpAs from these pathogens bound specifically to their cognate transferrins in our assay. We used a similar approach to evaluate whether *Hs*TbpB binding was specific for bovine transferrin. A displacement ELISA using soluble GST-*Hs*TbpB showed that unlike *Hs*TbpA, *Hs*TbpB binds bovine, ovine, and caprine transferrins, but has higher affinity for bovine transferrin (Figure 2C). Human transferrin was used as a negative control.

**Figure 2.**
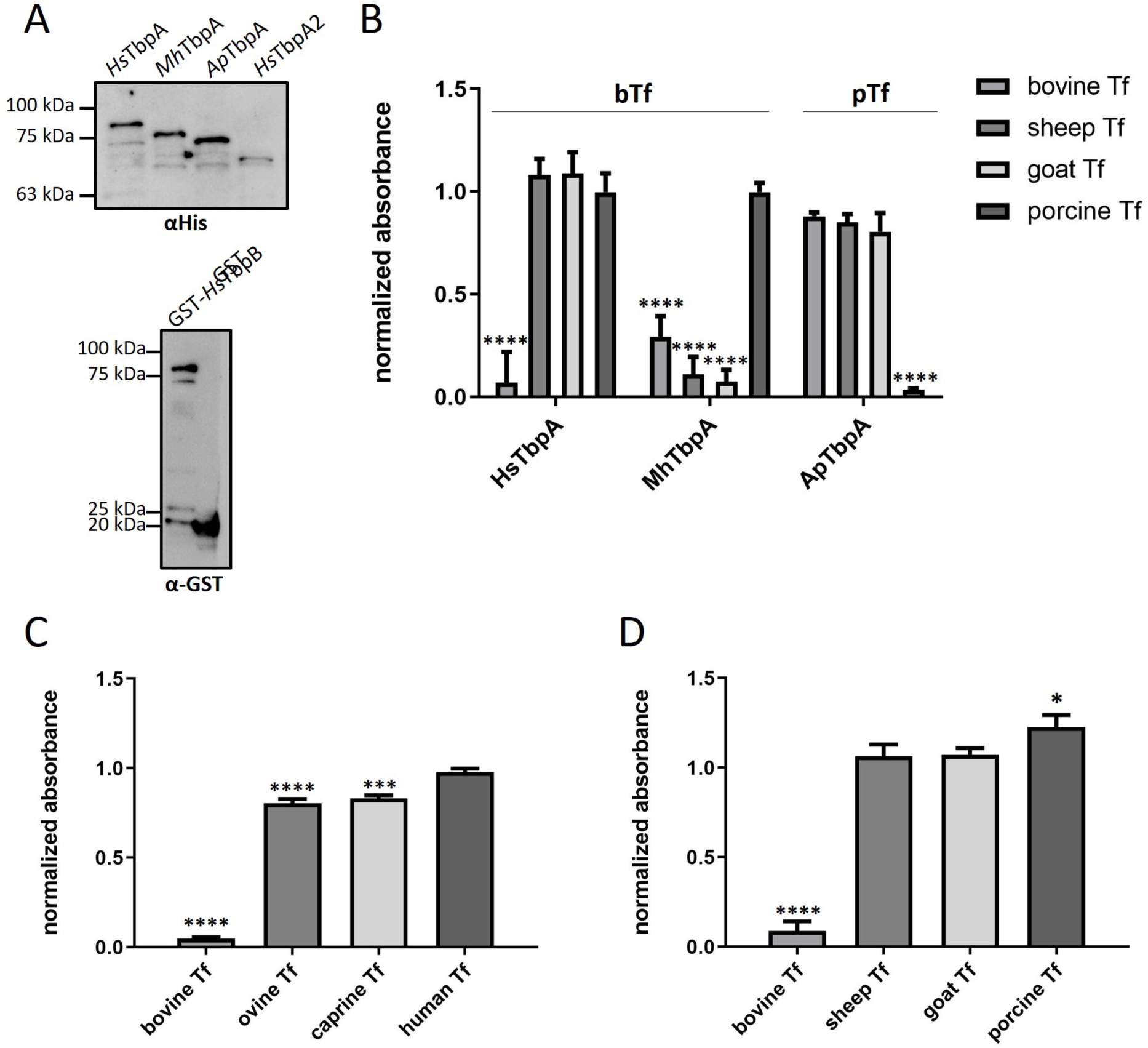
Displacement ELISAs with *Hs*TbpA, *Hs*TbpB, and *Hs*TbpA2 individually expressed in *E. coli*. (A) Anti-his (top) and anti-GST westerns showing expression of the TbpAs, TbpA2, and TbpB used in the ELISA assays. Displacement ELISAs were performed separately for TbpA (B), TbpB (C) and TbpA2 (D). ELISAs were performed and analyzed as described in Figure 23, with some modifications. Tbps were overexpressed in *E. coli*. Whole unfixed *E. coli* cells (B and D) or GST-tagged proteins in *E. coli* lysate (C) were immobilized on ELISA plates. Labeled transferrins (bovine for *Hs*TbpA, *Mh*TbpA, *Hs*TbpA2, and *Hs*TbpB, and porcine for *Ap*TbpA) were mixed with either buffer or a ∼40-fold excess of unlabeled competitor transferrin. The identity of the competitor transferrins is shown on the x-axis. See Methods sections 4.3.4 and 4.3.5 for more details. Results were analyzed by one-way ANOVA followed by Dunnett’s test, with all comparisons made to the “no competitor” column. Readings represent the mean of three independent experiments and error bars represent standard error of the mean.

We also asked whether our strain of *H. somni* encoded a TbpA2, and whether this receptor was as selective as TbpA. At the time these experiments were performed, a limited number of *H. somni* genomes were available in NCBI and it was unclear how widespread TbpA2 was among *H. somni* strains. We amplified out *Hs*TbpA2 from strain H191, and cloned and expressed it in *E. coli*. We then performed a displacement ELISA in the same way as had been performed for TbpA. *Hs*TbpA2 clearly bound bovine transferrin, which supports a previous report illustrating that the *H. somni* TbpA2 homolog is a transferrin binding protein. Moreover, *Hs*TbpA2 like, *Hs*TbpA, appears to selectively bind bovine transferrin in this assay (Figure 2D).

While our displacement ELISAs suggested that TbpB is responsible for the binding of *H. somni* to multiple bovid transferrins, we decided to complement our competitive binding experiments with direct binding assays to gain additional support for this result. We performed direct binding ELISAs by immobilizing transferrin on 96-well plates and detecting the binding of detergent-extracted *Hs*TbpA and *Hs*TbpA2 (Figure 3A, B). As in the displacement ELISA, detergent-extracted *Hs*TbpA bound bovine transferrin, and not ovine, caprine, or porcine transferrins (Figure 3A). *Hs*TbpA2, on the other hand, showed binding to bovine, ovine, and caprine transferrins, although the absorbance signal for the latter two interactions was only a quarter of what was observed for binding of *Hs*TbpA2 to bovine transferrin (Figure 3B).

**Figure 3.**
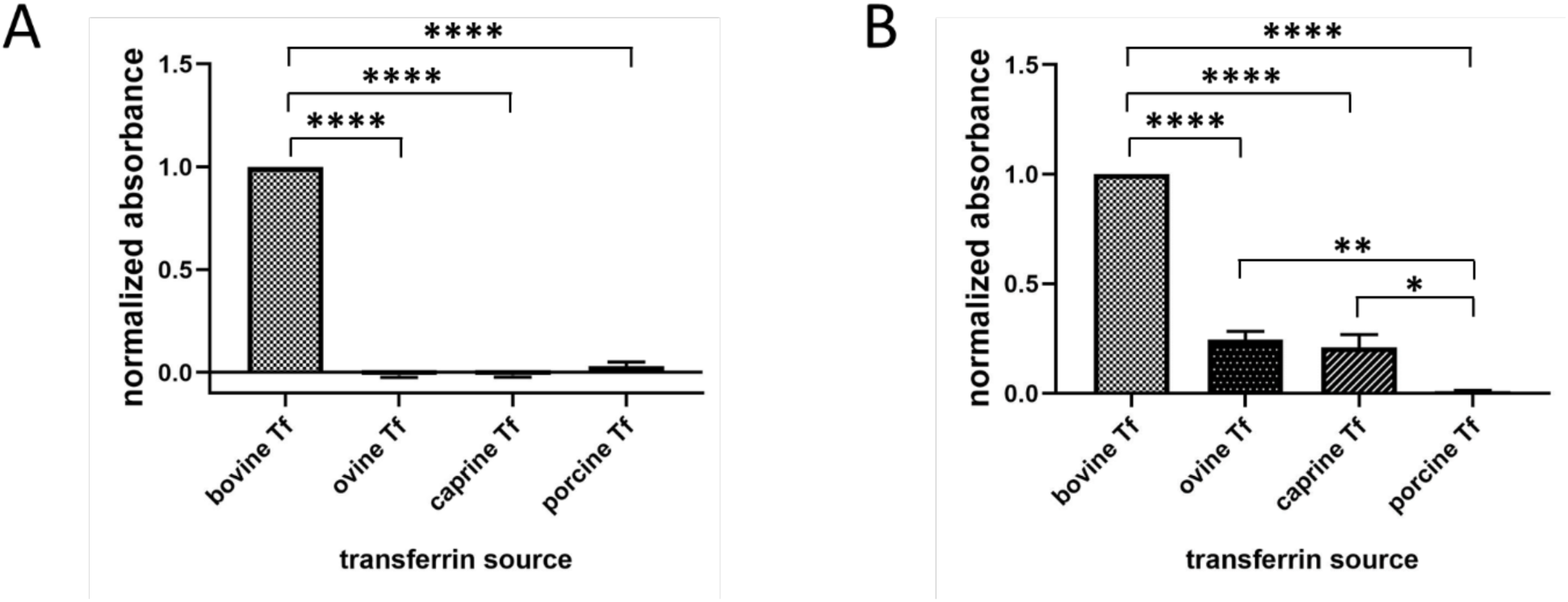
Direct binding assays with *Hs*TbpA and *Hs*TbpA2. ELISAs measuring binding between *Hs*TbpA (A) and *Hs*TbpA2 (B) and unlabeled bovine, ovine, caprine, and porcine transferrins. 1 µg of purified unlabeled transferrin was applied to each well. His-tagged TbpA and TbpA2 were expressed in *E. coli* and extracted using detergent. Binding was detected using anti-his antibodies. Readings were normalized to the signal obtained using bovine transferrin to account for total signal variation between experiments. Data was analyzed by one-way ANOVA followed by Tukey’s test. Results represent the means of three independent experiments and error bars represent standard error of the mean.

Because *Hs*TbpA2 binds to ovine and caprine transferrins and is a member of the TBDT family, we concluded that the growth observed in the presence of ovine and caprine transferrins must at least be partially due to this receptor. In contrast, *Hs*TbpB cannot act as a transporter, but may be acting to promote iron uptake from ovine and caprine transferrins as well. To determine whether the affinity of *Hs*TbpB for ovine and caprine transferrins would allow it to bind these proteins at physiological concentrations, we decided to measure binding affinities between the bovid transferrins and *Hs*TbpB.

### The affinities of *H. somni* TbpB for ovine and caprine transferrins permit binding at physiological concentrations

We measured dissociation constants (K_D_’s) for *Hs*TbpB-transferrin interactions to determine if binding occurs at physiological concentrations. We used bovine, ovine, and caprine transferrins and compared the corresponding dissociation constants with the concentration of transferrin that would be available in the host (Figure 4A-C, Table 7). While TbpB had a 3-fold lower affinity for ovine transferrin and a 15.7-fold lower affinity for caprine transferrin compared to its affinity for bovine transferrin, both dissociation constants are far lower than the concentration of transferrin found in the serum, which ranges between just over 15 µM to just under 90 µM in healthy cattle (see Discussion) (29, 30). We also measured binding of a TbpB from *Neisseria meningitidis*, a human pathogen, to determine the affinity of TbpBs for non-cognate transferrins. We observed no binding of *Nm*TbpB to ovine transferrin within the concentration range used in the experiment (Figure 4D). Taken together, our results suggest that *H. somni* TbpB is more promiscuous than *Hs*TbpA, and may, along with *Hs*TbpA2, contribute to iron uptake from ovine and caprine transferrins by *H. somni*.

**Figure 4.**
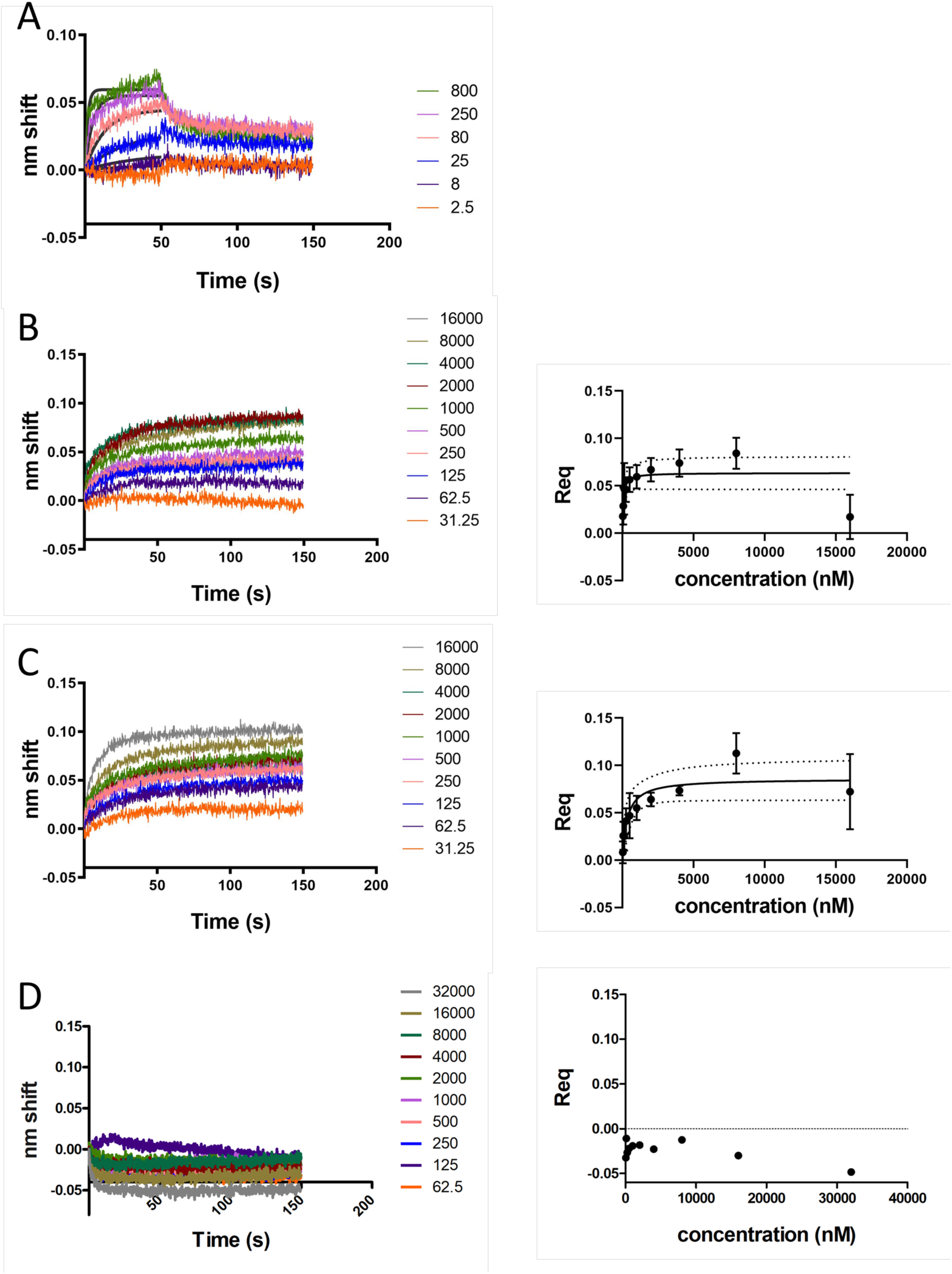
BLI sensorgrams and (when appropriate) saturation curves showing binding of *Hs*TbpB to ruminant transferrins. BLI was used to derive affinity constants for the interactions between *Hs*TbpB and the bovine, ovine, and caprine transferrins. Dissociation constant values are listed in Table 7. TbpB fused to a biotinylated BAP tag was immobilized on streptavidin sensors and binding was measured to varying concentrations of transferrin. Binding of *Hs*TbpB to bovine transferrin could be fit to a 1:1 binding model, but did not reach steady state (A). Conversely, the *Hs*TbpB interactions with ovine and caprine transferrins could not be fit to a 1:1 model and so a binding constant was derived by plotting saturation binding curves (B and C). The readings taken in the 135-145 second range were averaged and plotted versus concentration. Saturation curves were fit to a one site specific binding model. Dotted lines in B and C indicate 95% confidence intervals. Error bars represent the error of the mean of 3 independent experiments. Binding of *N. meningitidis* TbpB to ovine transferrin was also measured to ensure absence of nonspecific binding (D).

## Discussion

Many pathogens have host species-specific receptors that arise from a process of coevolution with the host. The sites that bear signatures of this coevolution tend to be functionally important, and so identifying them can lend insight into how a given receptor functions. Furthermore, host-specific interactions restrict pathogens to infecting a single host species or a narrow range of host species (31). This host specificity or host restriction can create a barrier to studying infection in animal models. A better understanding of the underlying mechanism of specificity could lead to the design of animal infection models that mimic the progress of disease more accurately(32).

The Tbps are host-specific receptors used by some host-restricted Proteobacteria for iron acquisition. The interaction between Tbps and their cognate transferrin has been shown to be a prerequisite for infection by Tbp-encoding bacteria. We looked for the most closely-related set of transferrins that could be distinguished by a Tbp system and found a paper reporting that *H. somni* binds bovine, but not ovine or caprine transferrins. Testing each receptor individually, we found that *H. somni* TbpA showed no detectable affinity for ovine or caprine transferrins.

However, we found the TbpB and an additional transferrin binding TBDT, TbpA2, did bind ovine and caprine transferrins. Furthermore, the affinities of *Hs*TbpB for the ovine and caprine transferrins is high enough to allow *Hs*TbpB to bind them in the blood. We observed that *H. somni* can utilize bovine, ovine, and caprine, but not human transferrin for growth. There are four possibilities that can account for our observations; firstly, *Hs*TbpA2 may work alone to promote iron uptake from ovine and caprine transferrins; alternatively, *Hs*TbpB may interact with TbpA2 to promote iron uptake; finally, *Hs*TbpB may work with *Hs*TbpA to promote iron uptake from the ovine and caprine transferrin C-lobes, with or without the accompaniment of *Hs*TbpA2, which is expected to take up iron from the N-lobe. A surface lipoprotein compensating for lack of function in a TBDT is a phenomenon that has been observed previously. For example, the function of a neisserial TbpA with mutations that reduced its binding to human transferrin could be partially rescued when it was co-expressed with TbpB (33). The mechanistic basis of this compensatory effect on the part of TbpB remains unknown. One possibility is that TbpB binding increases the local concentration of transferrin in the area around TbpA, thereby increasing the on rate enough to allow for iron uptake and thereby promoting growth. Another possibility is that the association of the transferrin-TbpB complex with TbpA leads to a conformational change that increases the affinity of TbpA for transferrin.

In this study, we conclude that TbpB can serve to broaden the host transferrin recognition range of *H. somni* because its affinity for ovine and caprine transferrins is high enough to promote binding in the blood. Our conclusions are based on reported serum transferrin concentrations. While there is a range of reported values, healthy cattle have transferrin concentrations of 1.37 to 6.6 g/L (29, 30). Using a molecular weight of ∼78 kDa for glycosylated bovine transferrin, the transferrin concentration in the blood can be estimated to range between 17 to 85 µM. This is similar to the serum transferrin concentrations found in other animals (34). All of the transferrins used in our experiments were fully iron loaded in order to remove a source of variability from the experiments. However, only about 25-50% of the transferrin found in the blood is iron saturated, meaning that in the blood, only ∼10-25% of the transferrin C-lobes are in the holo form (35). Because of this, the concentration ranges used in our experiments are approximately representative of the amount of transferrin that is available as an iron source in the blood for Tbp-containing bacteria. Another important consideration is that *H. somni* usually resides on mucosal surfaces where the transferrin concentration is significantly lower (although the exact concentration is difficult to measure). In human saliva, for example, the concentration of transferrin is roughly 10-50 nM (36–38). At these concentrations, the lower relative affinities of *Hs*TbpB and *Hs*TbpA2 for ovine and caprine transferrins compared to bovine transferrin may become significant. *H. somni* is isolated from cattle more frequently than from sheep or goats. It is possible that the concentration of transferrin on the mucosal surfaces of sheep and goats may be low enough to pose a barrier to *H. somni* colonization.

The *H. somni* transferrin receptors were chosen for specificity studies because they had been reported to distinguish between very similar transferrins. However, we did not find the *H. somni* Tbps to be as selective as reported. It is likely that the difference between our observations and previously published results is due to differences in assay sensitivity. Furthermore, because *Hs*TbpA2 is phase variable, it is possible that *Hs*TbpA2 was not expressed in the bacteria used in the whole-cell binding assays of the previously published study, which means that TbpA specificity may have had a larger relative effect on the results. It is worth noting that a previous study that identified TbpA2 in another strain of *H. somni* also suggested that *H. somni* has two sets of receptors, one specific for bovine transferrin and another that bound bovine, ovine, and caprine transferrins (16). However, this paper does not conclusively determine which Tbps are responsible for each binding pattern.

There were some discrepancies between the results obtained in the assays used in our study. Firstly, while *H. somni* appears to grow to similar levels on ovine and caprine transferrins, it appears to bind ovine transferrin with a higher affinity than caprine transferrin. This higher relative affinity for ovine transferrin compared to caprine was also observed in experiments using only *Hs*TbpA2 and only *Hs*TbpB. One possible explanation for this discrepancy is that growth-promoting synergistic interactions between Tbps were not picked up in the binding assays. Measuring the growth of *tbpB* and *tbpA2* knockouts on various transferrin iron sources would provide insight into whether and how these receptors work together. Secondly, binding of *Hs*TbpA2 to ovine and caprine transferrins was observed in the direct ELISA but not the displacement ELISA (Figure 24D and 25B). This is likely because the displacement ELISA did not have sufficient dynamic range to capture this difference in affinity. This difference would likely be observed at higher ratios of competitor to labeled transferrin. Interestingly, in the *Hs*TbpA2 displacement ELISA, the signal obtained using a competitor porcine transferrin was significantly higher than the signal obtained with no competitor (Figure 24D). This points to a flaw in the assay that cannot at present be explained. However, the difference in signal between the ovine and porcine, and caprine and porcine columns was not significant. It is clear that a more quantitative approach with purified TbpA2 is needed for follow-up binding assays. Measuring the binding affinities between *Hs*TbpA2 and the bovine, ovine, and caprine transferrins would provide the most accurate information about the relative strengths of these interactions.

We originally intended to mutate residues in ovine transferrin to the corresponding residues in bovine transferrin and screen for changes in affinity to the *H. somni* Tbps. This could still be done to identify important sites of contact between the Tbps and transferrin. However, in the case of *Hs*TbpB and *Hs*TbpA2, this would not be a screen for “all-or-none” changes in binding, but instead would likely involve precise measurements of changes in affinity. BLI, or a similarly sensitive technique that allows for quantitative binding analysis, would be ideal for these kinds of screens. Because of the presence of intermediate-affinity interactions, the *H. somni* Tbps may not be a good model system for studies looking at how to alter binding specificity in Tbp-transferrin interactions.

TBDT-dependent nutrient uptake systems can be either be monopartite, bipartite, or even multipartite, such as the Sus system in *Bacteroides* (39). TBDTs do not inherently require affiliated SLPs or other TBDTs to function. What then are the potential advantages of multipartite systems? TbpA is always found in an operon with TbpB and the latter has been shown to be essential for host colonization in the absence of lactoferrin receptors (12). However, the reason for this is not understood. TbpB is hypothesized to be important because it preferentially binds holo-transferrin. TbpA has no preference for transferrin’s iron loaded state, and so TbpB is thought to greatly increase the efficiency of iron acquisition. Furthermore, TbpA and TbpB have been shown to function synergistically and together, have a higher affinity for transferrin than either protein does alone. In this study we propose another advantage of having an accessory SLP: the expansion of substrate range. In multipartite nutrient uptake systems such as the Sus proteins, accessory lipoproteins are known to modulate substrate specificity (40). We propose that *Hs*TbpB also acts as a modulator, by broadening the specificity of the *H. somni* transferrin receptor.

To our knowledge, *H. somni* is the only pathogen that has three transferrin binding proteins. Whether *Hs*TbpA2 leads to the formation of a tripartite receptor is unknown, but the possibility is worth investigating. The three transferrin binding proteins in *H. somni* are reminiscent of the receptors HpuAB and HmbR in the pathogenic *Neisseria*. HpuAB forms a TbpBA-like bipartite receptor and binds hemoglobin-haptoglobin complexes, while HmbR is a single TBDT that binds hemoglobin (41, 42). Like the corresponding *H. somni* genes, both *hpuA* and *hmbR* are phase variable. Interestingly, the expression of HmbR has been correlated with a more virulent phenotype (43, 44). It is unknown whether TbpA2 increases virulence in *H. somni*. However, it is interesting to note that the commensal *H. somni* isolate 129PT (GenBank accession: CP000436) has a frame-shifted TbpA2 while more virulent isolates such as strain 2336 (CP000947) do not display this frameshift (unpublished observations). In the same way that it apparently benefits *Neisseria* to express three hemoglobin binding proteins, it may benefit *H. somni* to express three Tbps. In addition to the ability to bind both lobes of transferrin, we propose that having TbpBA and TbpA2 provides another advantage; *Hs*TbpB and *Hs*TbpA2 broaden transferrin recognition range to include ovine and caprine transferrins, and may play a role in allowing *H. somni* to infect these bovid species.

### d. Experimental Procedures

#### Plasmids and strains

*Histophilus somni* strain H191, an isolate collected from a healthy animal in a feedlot via nasal swab, was provided by Dr. Douglas W. Morck (University of Calgary) and was used for all *H. somni* growth and binding experiments. *Escherichia coli* C43(DE3) were used to express recombinant TbpAs and TbpBs. TbpBs were cloned into a custom vector designed by the Schryvers lab (45). TbpBs were cloned either into a custom vector designed by the Schryvers lab (University of Calgary) or into pGEX-6p-1. Constructs cloned into the custom vector contained an N-terminal biotin acceptor peptide (BAP), a 6x polyhistidine tag, maltose binding protein (MBP), and a Tobacco Etch Virus (TEV) protease cleavage site. Constructs expressed from pGEX-6p-1 contained an N-terminal GST tag followed by a PreScission Protease cleavage sequence. *Hs*TbpB (aa 21-643, where residue 1 is Cys1 of the mature peptide) was expressed cytoplasmically without its anchor peptide. TbpAs were provided by the Schryvers lab and were re-cloned into the pET26b+ vector and expressed with a PelB signal peptide, and an N-terminal 10x polyhistidine tag followed by a TEV cleavage site. (The C-terminal polyhistidine tag encoded in the vector was not used.)

#### Transferrin purification

Bovine holo-transferrin was purchased from Sigma. Ovine and caprine transferrins were purified from sheep and goat sera (Sigma). 50-100 mL of serum was used per purification. The serum samples were centrifuged at 3000*g* to remove insoluble components and initially fractionated via ammonium sulfate precipitation. Ammonium sulfate was added to the serum to achieve a 40% saturation at 4°C and dissolved at room temperature by end-over-end mixing. Once no more undissolved ammonium sulfate crystals were visible in the serum, the serum was centrifuged at 12 000*g* to remove precipitate. It was often necessary to perform two centrifugation steps, transferring the soluble fraction to a new tube in between steps, to remove all the precipitate. Further ammonium sulfate cuts were performed by adding ammonium sulfate to saturations of 55%, 70%, and 85% at 4°C. The 70% fraction, which is expected to contain most of the transferrin, was resuspended in 50 mM Tris, pH 8.0, and dialyzed in 2 L of the same buffer for 8-16 hours at 4 C, followed by 3 more changes of 2 L. Samples of the other fractions were dialyzed concurrently and used for SDS-PAGE analysis.

The dialyzed 70% ammonium sulfate fraction was further purified using anion exchange chromatography using an in-house packed 25 mL Q-Sepharose column. After multiple rounds of chromatography, primarily done to separate transferrin from albumin, the protein was concentrated and size exclusion chromatography using an S75 column (GE) was used for buffer exchange into PBS. The protein was concentrated further (if necessary), aliquoted, flash-frozen, and stored at −80 C.

#### Whole cell ELISAs with *H. somni*

*H. somni* strain H191 was grown overnight at 37°C and 5% CO2 on chocolate agar plates containing 1% IsoVitalex. Cells were harvested from plates, washed once in PBS, and resuspended at an OD_600_ of 0.05 in BHI broth containing 0.5% yeast extract, 0.05% thiamine pyrophosphate and 50 uM deferoxamine, and left to grow for 16 hours overnight at 37°C with shaking. Cells were harvested by centrifugation for 10 minutes at 4000*g*, washed once in PBS containing 10 mM MgCl_2_, and fixed for 20 minutes in 1.85% formaldehyde solution (3.7% formaldehyde mixed with equal parts PBS + 10 mM MgCl_2_). Cells were washed once in PBS + 10 mM MgCl_2_ and resuspended at an OD_600_ of 10 in the same buffer. 100 uL of cells per well was applied to 96-well flat-bottomed tissue culture plates (Sarstedt). Plates were incubated for 3 hours at room temperature in a biosafety cabinet. Plates were then washed once with PBS and patted dry. 50 uL of 6.6 nM biotinylated transferrin and 50 uL of 132 nM of unlabeled transferrin was applied to each well and left to incubate for 2 hours at room temperature. Transferrin was diluted in 5% skim milk dissolved in PBS. Plates were washed 3 times with PBS and 100 uL volumes of streptavidin-HRP at a dilution of 1 in 5000 dissolved in skim milk solution was added to the wells and left to incubate for one hour at room temperature. Plates were washed 3 times with PBS. Binding was detected with 3,3’,5,5’-Tetramethylbenzidine (TMB) substrate (Sigma), reactions were stopped with 2 M HCl, and absorbance was read at 450 nm. Signal of wells containing buffer instead of cells was subtracted from all reads. Wells including cells but not labeled transferrins were also used to monitor for nonspecific binding. Absorbance readings were normalized to the signal of wells containing no competitor. Data was plotted and analyzed in Graphpad Prism. Statistical analysis was performed using one-way ANOVA followed by Dunnett’s post test. Normalized readings were compared to a value of 1, the normalized signal for the “no competitor” column, and the value that is expected when no binding occurs.

#### Whole-cell ELISAs with *E. coli*

5 colonies each of *E. coli* C43 were inoculated into 1 mL of LB medium containing 5 ug/mL kanamycin sulfate. The starter culture was grown for 16 hours overnight with shaking at 37C. The starter culture was used to inoculate TB medium containing 50 ug/mL of kanamycin sulfate. The dilution was 1 in 100. Cells were grown for approximately 1.5 hours to an OD_600_ of 0.15-0.2 and induced with 0.5 mM isopropyl β-D-1-thiogalactopyranoside (IPTG). Cells were grown for another 1-1.5 hours to an OD_600_ of ∼0.3, harvested, and resuspended in PBS to a density of 10^10^ cells/mL. Expression was verified by immunoblotting with anti-his antibodies (dilution of 1 in 5000) (Thermo Fisher Scientific Cat# MA1-21315-HRP, RRID:AB_2536989). 100 uL volumes of cell suspension were added to 96-well tissue cultures plates (Sarstedt) and allowed to adhere at 37° C for 3 hours. Plates were washed once with PBS and blocked for 1 hour with 5% skim milk solution. The rest of the procedure was carried out as described for the *H. somni* ELISA, except that the concentration of transferrin used was 250 nM instead of 132 nM (∼38-fold excess of unlabeled competitor). Data was analyzed as described in 4.3.4.

#### ELISAs with *E. coli* lysates containing GST-TbpB

*E. coli* C43 starter cultures were used to inoculate 5 mL of 2YT-based ZYP-5052 autoinduction media containing 100 µg/mL of ampicillin. Cultures were left to grow for 48 hours at 20°C. The low temperature was used to minimize formation of inclusion bodies. Cultures were harvest by centrifugation at 10000*g* and resuspended in 1 mL of PBS pH 7.4, 1 mg/mL lysozyme, 1 mM DnaseI, 1 mM benzamidine. Cells were lysed by sonication (30 seconds of constant pulsing at the lowest setting using a microtip followed by 1 minute of incubation on ice, repeated 1-2 more times). Insoluble components were pelleted by centrifugation at 16000*g* for 10 minutes at 4°C. Expression was verified by immunoblotting with anti-GST antibodies (dilution 1 in 5000) (GenScript Cat# A00867, RRID:AB_982273). Lysates were applied to ELISA plates (96-well tissue culture plates from Sarstedt or 96-well Nunc MaxiSorp plates from Invitrogen) and left to incubate overnight at 4°C. Plates were washed 3 times with 100 µL volumes of PBS-T. Bovine transferrin was labeled with HRP using an EZ-Link ™ Plus Activated Peroxidase Kit (ThermoFisher). Mixtures of HRP-labeled bovine transferrin (final concentration of 6.6 nM) and unlabeled bovine, ovine, caprine, or human transferrins (final concentration of 250 nM, ∼38-fold excess of unlabeled competitor) were added to the plates (100 µL/well) and left to incubate for 1 hour at room temperature. Plates were washed 3 times with PBS-T. Signal read-out was performed by adding 100 µL of 3,3’,5,5’-Tetramethylbenzidine (TMB) substrate (ThermoFisher). The reaction was stopped by addition of 100 µL of 2 M HCl and absorbance at 450 nm was read using an Epoch Microplate Spectrophotometer (BioTek). Signal of wells containing GST alone instead of GST-TbpB was subtracted from all reads. Wells including cells but not labeled transferrins were also used to monitor for nonspecific binding. Absorbance readings were normalized to the signal of wells containing no competitor. Data was analyzed as described in 4.3.4.

#### ELISAs with whole-cell extracts containing His-tagged TbpAs

100 µL volumes of purified transferrin at a concentration of 10 µg/mL were applied to either 96-well tissue culture plates (Sarstedt) or to Nunc MaxiSorp ™ ELISA plates (Invitrogen). Plates were incubated overnight at 4°C then washed 3 times with PBS-T. TbpAs were expressed in *E. coli* as described in section 4.3.4. Cells were pelleted and resuspended by slow pipetting in one-fifth the culture volume in PBS pH 7.4 containing 3% Elugent ™ (Millipore), 1 mg/mL lysozyme, 1 mM DnaseI and 1 mM benzamidine. Final extract volumes ranged from 5 to 10 mL. Extractions were performed by slow stirring for one hour at room temperature or overnight at 4°C. Following extraction, undissolved components were removed by centrifugation at 16000*g* for 10 minutes at 4°C. 100 µL of extract was applied to transferrin-coated wells and plates were incubated for 1 hours at room temperature. Wells were washed 3 times with buffer containing 1% Elugent. Wells were then incubated, either for 1 hour at room temperature or overnight at 4°C, with 100 µL of mouse anti-his antibody (ThermoFisher) diluted 1 in 5000 in PBS containing 5% skim milk and 1% Elugent. Wells were washed 3 times with buffer containing 1% Elugent then incubated for 1 hour at room temperature with 100 µL of goat anti-mouse antibody in PBS containing 5% skim milk, 1% Elugent. Wells were washed 3 times with buffer containing 1% Elugent. Signal read-out was performed by adding 100 µL of 3,3’,5,5’-Tetramethylbenzidine (TMB) substrate (ThermoFisher). The reaction was stopped by addition of 100 µL of 2 M HCl and absorbance at 450 nm was read using an Epoch Microplate Spectrophotometer (BioTek).Readings were normalized to the absorbance reading obtained with bovine transferrin. This was done to account for variations in signal observed between experiments. These variations were due to slight differences in protein concentration, adhesion to the plates, and incubation times with TMB. Data was analyzed by one-way ANOVA followed by Tukey’s test.

#### Lysate preparation for binding experiments involving in vivo biotinylated TbpBs

BAP-tagged TbpBs were expressed from *E. coli* C43(DE3) cells. Five colonies per construct were inoculated into 2YT-autoinduction media (45). Because expression levels varied between TbpB orthologs, different culture volumes were used to produce each construct: 10 mL for *Hs*TbpB, 5 mL for *Mh*TbpB, and 2 mL for *Nm*TbpB. Cells were grown overnight for 20 hours with shaking at 37°C. Cells were pelleted at 10 000*g* for 5 minutes at 4°C, resuspended in 1 mL of PBS containing 1 mg/mL lysozyme and 1 mM PMSF, and lysed by sonication. Cell debris was removed by centrifugation at 16 000*g* for 10 minutes at 4°C.

#### Bio-layer interferometry binding assays

Bio-layer interferometry was performed using the Octet RED96 system (PALL-ForteBio). All samples were diluted in kinetics buffer (1x PBS, 0.1 mg/mL bovine serum albumin, 0.002% Tween-20). Streptavidin-coated sensors were first equilibrated in kinetics buffer, then dipped into lysates diluted twofold in kinetics buffer. After 100 seconds of loading, the sensors were washed for 200 seconds in kinetics buffer and then dipped into wells containing transferrin diluted in kinetics buffer. After 60 seconds of association, the sensors were dipped into buffer wells for 200 seconds to allow dissociation to occur. Sensors were regenerated in buffer consisting of 50 mM citrate, pH 4.5, and 100 mM EDTA. The wash-association-dissociation-regeneration sequence was repeated for every concentration of transferrin on the BLI plate.

#### *H. somni* growth experiments

*H. somni* was grown overnight at 37 °C on chocolate agar plates. Cells were swabbed from plates, resuspended in BHI media containing 0.5% yeast extract and 0.1% Tris (BHI-YT), harvested by centrifugation and then resuspended again in BHI-YT. 1 mL of BHI-YT containing 0.01% thiamine pyrophosphate and 10 µM of partially iron-saturated bovine transferrin was inoculated with the bacterial suspension at an OD_600_ of 0.1. Cultures were left to grow at 37 °C with shaking at 180 rpm in capped and parafilm-sealed 15 mL conical tubes. After 6 hours of growth, cells were transferred to a 2 mL tube, harvested by centrifugation, washed once in 1X PBS pH 7.4, and resuspended again in the same buffer. The transferrin-dependent growth experiment was performed in BHI-YT containing 0.01% thiamine pyrophosphate, 1 mM deferoxamine, and either PBS or a transferrin iron source. Iron sources consisted of iron-loaded bovine, ovine, caprine, or human transferrin in PBS at concentrations of 25, 12.5, 6.25, and 3.125 µM. *H. somni* was inoculated at an OD_600_ of 0.02 and left to grow overnight without shaking at 37 °C, 5% CO_2_. The growth was performed in 50 µL volumes in flat-bottomed tissue culture plates (Sarstedt) which were covered with an air-permeable seal (SigmaAldrich). After 20 hours of growth, OD_600_ was measured using a Victor Nivo plate reader (Perkin-Elmer) at 37 °C following 1 minute of shaking. 10 µL of culture from the 25 µM transferrin condition was used to measure cfu/mL. Serial dilutions of these samples were plated on chocolate agar plates and colony forming units were counted following another 16 hours of growth.

## Acknowledgements

We would like to thank members of the Moraes and Schryvers lab for their feedback on the manuscript. We would also like to thank Dr. Anthony Schryvers for providing plasmids and strains. This work was supported by a grant from the Natural Sciences and Engineering Research Council of Canada (NSERC, RGPIN-2018-06546), Canada Foundation for Innovation (CFI) and the Ontario Ministry of Economic Development and Innovation (MEDI). TFM is supported by a CRC in the Structural Biology of Membrane Proteins.

---

**Table 1.** *Hs*TbpB binding constants for interactions with bovine, ovine, and caprine transferrins. Binding constants were derived by fitting to a 1:1 binding model (for bovine transferrin) or by plotting readings at steady state and fitting the saturation binding curves. All values are means of three separate experiments +/- standard error.

| | $k_{on}$ (nM<sup>-1</sup>s<sup>-1</sup>) | $k_{off}$ (s<sup>-1</sup>) | $K_D$ (nM) |
| --- | --- | --- | --- |
| Bovine Tf | 916000 +/- 531000 | 0.0403 +/- 0.0219 | 21.7 +/- 6.9 |
| Ovine Tf | n.a. | n.a. | 65.2 +/- 51.2 |
| Caprine Tf | n.a. | n.a. | 340 +/- 194 |

